# Fluorescence-based detection of natural transformation in drug resistant *Acinetobacter baumannii*

**DOI:** 10.1101/262311

**Authors:** Anne-Sophie Godeux, Agnese Lupo, Marisa Haenni, Simon Guette-Marquet, Gottfried Wilharm, Maria-Halima Laaberki, Xavier Charpentier

## Abstract

*Acinetobacter baumannii* is a nosocomial agent with a high propensity for developing resistance to antibiotics. This ability relies on horizontal gene transfer mechanisms occurring in the *Acinetobacter* genus, including natural transformation. To study natural transformation in bacteria, the most prevalent method uses selection for the acquisition of an antibiotic resistance marker in a target chromosomal locus by the recipient cell. Most clinical isolates of *A. baumannii* are resistant to multiple antibiotics limiting the use of such selection-based method. Here we report the development of a phenotypic and selection-free method based on flow cytometry to detect transformation events in multidrug resistant (MDR) clinical *A. baumannii* isolates. To this end, we engineered a translational fusion between the abundant and conserved *A. baumannii* nucleoprotein (HU) and the superfolder green fluorescent protein (sfGFP). The new method was benchmarked against the conventional antibiotic selection-based method. Using this new method, we investigated several parameters affecting transformation efficiencies and identified conditions of transformability one hundred times higher than those previously reported. Using optimized transformation conditions, we probed natural transformation in a set of MDR clinical and non-clinical animal *A. baumannii* isolates. Regardless of their origin, the majority of the isolates displayed natural transformability, indicative of a conserved trait in the species. Overall, this new method and optimized protocol will greatly facilitate the study of natural transformation in the opportunistic pathogen *A. baumannii*.

## IMPORTANCE

Antibiotic resistance is a pressing global health concern with the rise of multiple and pan-resistant pathogens. The rapid and unfailing resistance to multiple antibiotics of the nosocomial agent *Acinetobacter baumannii*, notably to carbapenems, urges to understand how it acquires new antibiotic resistance genes. Natural transformation, one of horizontal gene transfer mechanisms in Bacteria, was only recently described in *A. baumannii* and could explain its ability to acquire resistance genes. We developed a reliable method to probe and study natural transformation mechanism in *A. baumannii*. More broadly, this new method based on flow cytometry will allow experimental detection and quantification of horizontal gene transfer events in multidrug resistant *A. baumannii.*

## INTRODUCTION

*Acinetobacter baumannii* is a gram-negative bacterium responsible for healthcare associated infections in humans and animals (1, 2). Outside the hospital, it has been detected in various environments (3) but its exact reservoir remains unclear. Asymptomatic carriage of *A. baumannii* has been reported in human and animals although the prevalence in healthy individual seems low (4–7). Although rarely encountered in animals, *Acinetobacter* infections represent a major challenge for physicians in intensive care units as acquired antibiotic resistance is widespread in *A. baumannii* isolates. Worldwide, the percentage of invasive isolates with combined resistance to fluoroquinolones, aminoglycosides and carbapenems is on the rise (8, 9). The ability of this bacterium to acquire multiple antibiotic resistance genes is well established but the underlying mechanisms are not clearly understood. Genome sequencing of multidrug-resistant (MDR) strains revealed that multiple events of horizontal gene transfer are responsible for the acquired multidrug resistance (10, 11) and that extensive recombination events drive the diversification of *A. baumannii* resulting also in antigenic variation (12). Among the horizontal gene transfer mechanisms, natural transformation has been demonstrated in some *A. baumannii* isolates, providing a plausible route for intra- and interspecific genetic exchanges (13–15). Natural transformation could notably explain the frequent occurrence in *A. baumannii* genomes of large genomic islands lacking features characteristic of self-transmissible elements, such as the AbaR genomic islands (16, 17). These islands carry multiple resistance genes and have also been revealed in pathogenic non-*baumannii* species (18). Natural transformation allows a bacterial cell to take up exogenous DNA and subsequently incorporate it into its genome through homologous recombination (19). Although some sequence identity is required between the donor DNA and the recipient’s chromosome for recombination, natural transformation allows integration of more diverse sequences such as transposons, integrons and gene cassettes from distant species (20). The recombination of exogenous DNA into the chromosome requires that the bacteria first enter the physiological state of competence which includes the expression of a molecular machinery to take up DNA, protect it from degradation and bring to the chromosome. Although natural transformation is a conserved trait in bacteria, the conditions required to trigger competence and to transform are often elusive and species-specific (19, 21). Interestingly, some antibiotics were found to induce competence in three distinct human pathogens – namely *Helicobacter pylori, Legionella pneumophila* and *Streptococcus pneumoniae*, raising the concern that some antibiotic treatment may increase horizontal gene transfer events (22–24). More specifically in *A. baumannii* and related species, description of natural transformation is rather recent and occurs upon movement on wet surfaces (13, 15).

Natural genetic transformation is generally detected through a phenotypic outcome. Acquisition of antibiotic resistance conferred to the cells by the chromosomal integration of a provided DNA fragment carrying an antibiotic resistance gene represents the most common phenotypic outcome (13). However, as most clinical isolates of *A. baumannii* are resistant to multiple antibiotics, using antibiotic selection to evaluate natural transformation in *A. baumannii* appears both experimentally challenging and ethically questionable. Thus, an alternative method to detect and quantify transformation events is needed to explore natural transformation in MDR clinical *A. baumannii* isolates.

Here we present a phenotypic and selection-free method based on flow cytometry to detect transformation. To this end, we developed a flow cytometry-optimized fluorescent marker consisting of a chromosomal translational protein fusion between a nucleoid associated protein (HU) and the superfolder GFP (sfGFP). Flow cytometry effectively discriminates HU-sfGFP green fluorescent *A. baumannii* cells among a large population of non-fluorescent cells. We demonstrated that DNA encoding the HU-sfGFP marker is a suitable substrate for genetic transformation in *A. baumannii*. Compared to selection-based detection using an antibiotic resistance marker, flow cytometry combined with the HU-sfGFP marker offers a more reliable and direct quantification of transformants. We took advantage of this transformation detection method to improve transformation conditions for *A. baumannii* and to probe natural transformation in MDR clinical isolates but also in non-clinical strains.

## RESULTS

### Optimization of a bright chromosomally-encoded GFP fusion in *A. baumannii*

We sought to use expression of GFP as a fluorescence-based phenotypic outcome of natural transformation. To allow distinction of bacterial transformants from a larger non-fluorescent bacterial population, an optimal fluorescent marker should therefore confer a fluorescence bright enough to be discriminated from bacterial autofluorescence. Moreover, in order to be a suitable substrate for natural transformation in various *A. baumannii* isolates, this marker must insert in a conserved locus in the *A. baumannii* chromosome. To select for an optimal fluorescent marker, we engineered several GFP translational fusions to abundant *A. baumannii* proteins in order to increase fluorescence of bacterial cells in comparison to cytoplasmic (free) GFP. We first took advantage of the recent strategy developed by Kjos *et al*. that engineered a bright strain of *S. pneumoniae* using a translational C-terminal fusion of the superfolder GFP (sfGFP) to HlpA, the unique nucleoid-associated protein of this gram-positive pathogen (25). In gram-negative bacteria, twelve nucleoid-associated proteins (NAP) are commonly described (26). Using the BLAST algorithm, we searched for genes encoding nucleoid-associated proteins in the genome of the multidrug-resistant clinical isolate *A. baumannii* AB5075. We identified three homodimeric small NAPs: HU (ABUW_2198), HNS (ABUW_3609) and Fis (ABUW_1533). Two other candidate proteins were also selected based on their abundance inferred from proteomic analysis performed in *A. baumannii* strain ATCC 17978 (27). In this particular strain, heat shock protein DnaK (ABUW_3879) and ribosomal protein S1 RpsA (ABUW_2242) were among the more abundant cytoplasmic proteins. Moreover, these two proteins appeared suitable candidates for GFP translation fusions based on successful C-terminal fusions in *Mycobacterium smegmatis* (28) and *Bacillus subtilis* (29). Consequently, we generated translational C-terminal fusions of the sfGFP with HU, HNS, Fis, DnaK and RpsA proteins. Lastly, as mean of comparison, we constructed a strain carrying the sfGFP-encoding gene (*sfgfp*) under a strong constitutive synthetic promoter inserted into a neutral chromosomic locus in *A. baumannii*. The chosen locus was the sulfite reductase gene (ABUW_0643 or *cysI*) identified previously as a conserved but non-essential gene in *A. baumannii* (13). The six genetic constructs were independently integrated into the chromosome of the pathogenic AB5075 *A. baumannii* strain (Fig. 1A.). *A. baumannii* was also transformed with a broad host-range and multicopy plasmid carrying the *sfgfp* gene under a strong constitutive promoter (plasmid pASG-1). Using western blot analysis of bacterial lysates, we confirmed that all the AB5075 derivative strains expressed sfGFP protein fusions with the expected molecular weight (Fig. S1). However, the Fis-sfGFP appeared less expressed than other sfGFP protein fusions.

**Figure 1.**
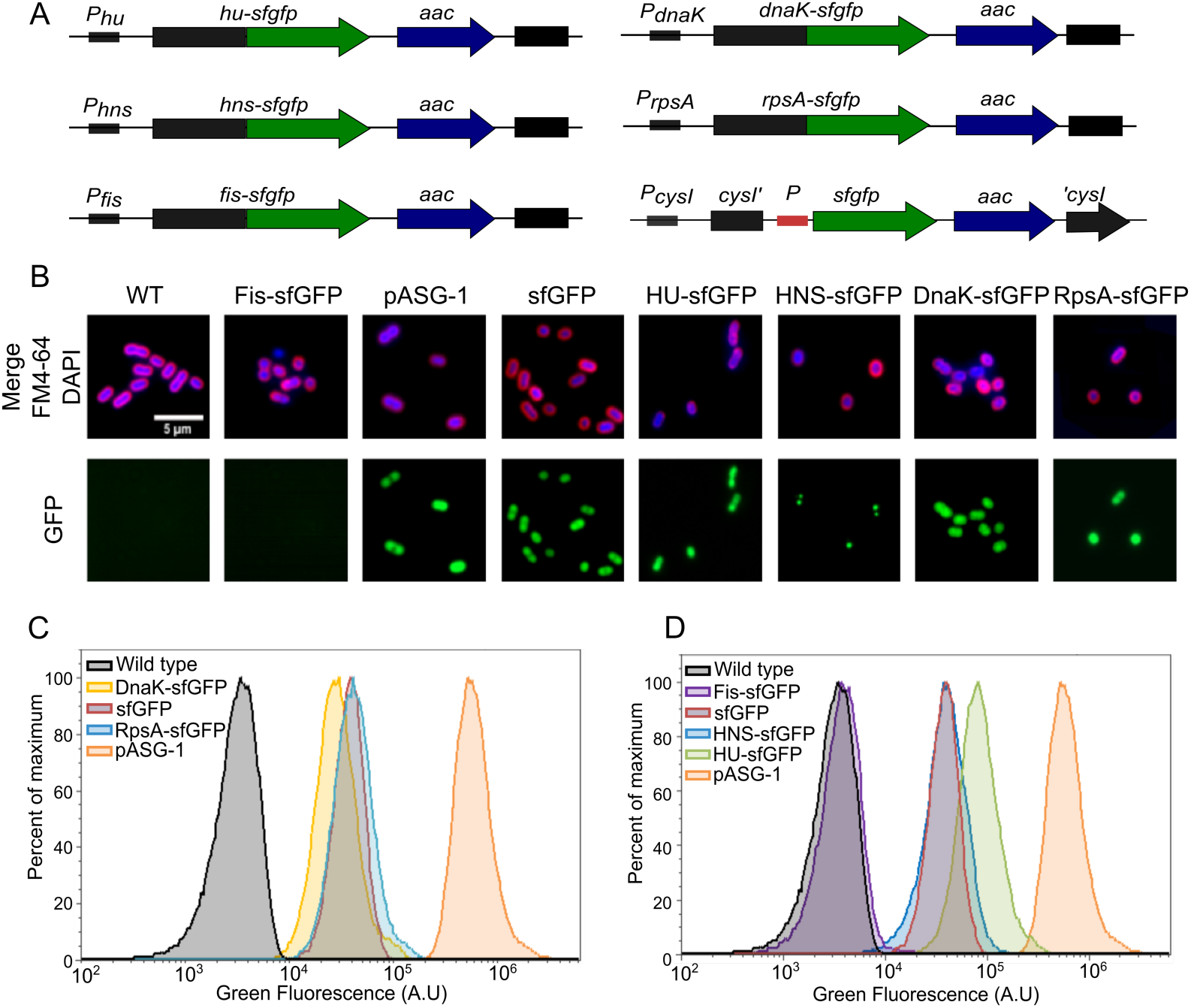
Engineering a chromosomal fluorescent marker in *A. baumannii.* A. Schematic representation of genetic constructions encoding the fluorescent markers used in this study. In green is represented the gene encoding for the sfGFP, sfGFP fused to the C-terminus of HU, HNS, Fis, DnaK, RpsA proteins. In blue, an aac(3)IV gene encoding for resistance to apramycin. The red box corresponds to a synthetic constitutive promoter. The sequences and genes in black are part of the AB5075 chromosome. B. Fluorescence microscopy images of *A. baumannii* AB5075 derivatives expressing free or chimeric sfGFP proteins. Membranes were stained with FM4-64 and DNA with DAPI (upper row and first column). For illustration purpose, the green fluorescence intensities were normalized between the samples (lower row). C. Flow cytometry histograms of green fluorescence intensities of AB5075 derivatives expressing free sfGFP or sfGFP protein fusions in comparison to wild-type level (black curve). A representative experiment is shown.

Then, fluorescence signals and subcellular localizations of the various sfGFP protein fusions were analyzed using fluorescence microscopy (Fig. 1B.). Consistent with the immunoblotting analysis, the strain expressing the Fis-sfGFP marker presented a low green fluorescence that was comparable to the wild-type strain (autofluorescence, Fig. 1B). Otherwise, all other sfGFP fusions were detectable by fluorescence microscopy. Still, the cellular localization of the green fluorescence signal differs among the strains. As expected, free sfGFP expressed from either a chromosomal locus or from a plasmid (*cysI*::*sfgfp* or pASG-1) presented a diffuse localization pattern in bacterial cells. This localization pattern was similar for strains expressing sfGFP translation fusions with cytoplasmic proteins (DnaK-sfGFP and RpsA-sfGFP). In contrast, when sfGFP was fused to HU-sfGFP and H-NS-sfGFP, the GFP signal showed a discrete pattern. The HU-sfGFP signal appeared to co-localize with the nucleoid visualized by DAPI staining. Also, the subcellular localization of HNS-sfGFP was restricted to few discrete foci per cell.

Subsequently, green fluorescence intensities of the various strains were then quantified by flow cytometry (Fig. 1C.). All strains expressing sfGFP chromosomal markers, but Fis-sfGFP, exhibited a fluorescence distinct from the unmarked wild-type strain (autofluorescence). As expected, cytoplasmic sfGFP expressed from a replicative plasmid (pASG1) displayed a strong green fluorescence in comparison to the wild-type strain (200-fold when comparing the mean intensities) whereas the cytoplasmic sfGFP expressed from the chromosome presented an intermediate green fluorescence level (about 12-fold increase relative to autofluorescence). Among the chromosomal markers, the HU-sfGFP construct presented the strongest fluorescence with a mean of 20-fold increase compared to wild-type levels. The other genetic constructs (RpsA-sfGFP, DnaK and HNS-sfGFP) presented fluorescence levels comparable to the free sfGFP expressed from the *cysI* locus. Remarkably, the distribution of the fluorescence intensity of population expressing HU-sfGFP protein fusion does not overlap with the wild-type autofluorescence allowing a clear resolution of the two populations (Fig. 1C.). We ensured that the various fluorescent strains presented a growth similar to the wild-type strain indicating that both the translational fusions and their chromosomal insertions could be used for downstream applications (Fig. S2A.). Moreover, all fluorescent markers but Fis-sfGFP appear to be steadily expressed during growth in liquid medium with, expectedly, HU-sfGFP presenting the strongest signal among the six chromosomal constructions (Fig. S2B). In conclusion, among the six chromosomal fluorescent markers investigated, expression of the HU-sfGFP marker confers the highest fluorescence level and its expression is compatible with bacterial growth, both characteristics required to constitute a genetic marker of natural transformation.

### Benchmarking the HU-sfGFP chromosomal marker in *A. baumannii* transformation assay

In an *A. baumannii* population, the frequency of transformants does not usually exceed 10^-3^, meaning that less than 1 out of 1000 bacterial cells integrate and express a selection marker (13). Thus, a fluorescence-based phenotypic outcome of natural transformation must be able to detect a rare population of sfGFP-expressing cells within a large population of non-fluorescent cells. Because of its high-resolution power, antibiotic resistance selection-based assays are frequently used to detect rare transformation events. We thus benchmarked the direct flow cytometry-based detection of the HU-sfGFP marker to the selection-based antibiotic marker detection classically used for testing natural transformation. To this end, the HU-sfGFP marker bears an apramycin resistance gene (*aac(3)-IV*) allowing comparison of both methods of selection/detection for the same chromosomal locus of recombination (Fig. 1A). First, we tested the sensitivity and specificity of flow cytometry-based detection of bacterial cells expressing the HU-sfGFP marker within large non-fluorescent bacterial populations and compared it to the apramycin resistance selection. We subjected to both methods of detection the same samples consisting of serial dilutions of HU-sfGFP cells into wild type cells (Fig. 2).

**Figure 2.**
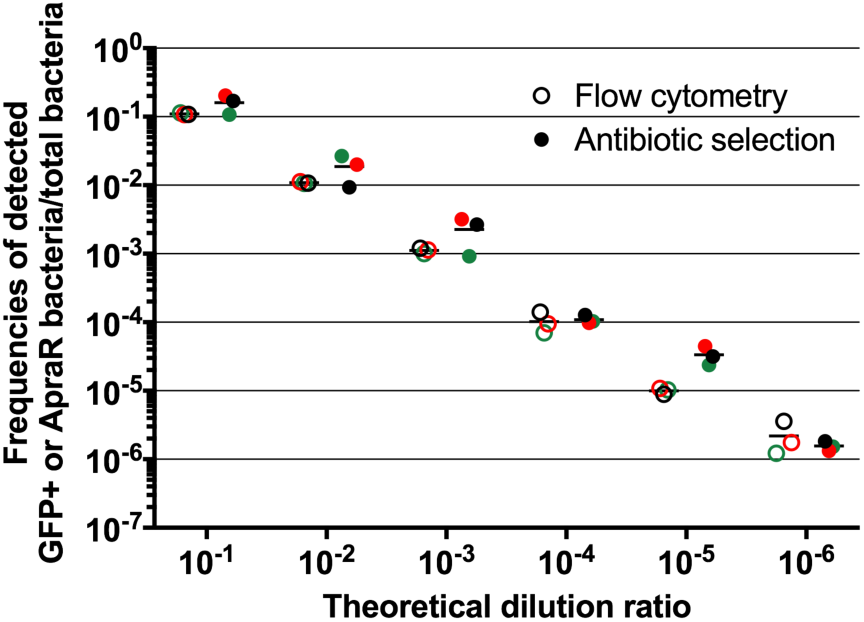
Comparison of paired measurements of hu-sfgfp_aac marker detection obtained by flow cytometry and antibiotic selection methods. HU-sfGFP cells were diluted into a suspension of wild-type *A. baumannii* AB5075 cells with increasing dilution factor (abscissa). Green fluorescent and apramycin resistant populations were determined (ordinate) both by flow cytometry (empty dots) and antibiotic selection (full dots). The three dots represent technical replicates. For a given dilution ratio, dots of the same color indicate a sample tested by both methods, the horizontal lines represent the mean of the three measures.

Both methods gave comparable determination of bacterial concentrations in the range of 10^-1^ to 10^-6^ dilution ratios and corresponded to the theoretical dilution factors. The latter dilution (10^-6^) represents the lower detection limit mostly for technical reasons. With our current equipment and settings, analysis of over 10^6^ bacterial cells is possible but would take at least five hours per sample.

We then tested the sensitivity and specificity of flow cytometry for detection of HU-sfGFP transformants within wild-type cells. To this end, we used the genomic DNA (gDNA) extracted from the *hu-sfgfp_aac* strain as substrate for natural transformation of wild-type AB5075 strain and compared the frequency of transformants measured either by flow cytometry or by antibiotic selection. In *Acinetobacter baylyi*, a non-pathogenic species, transformation efficiency increases in a DNA concentration-dependent manner (30). To assess the accuracy and reliability of the cytometry-based method, we used increasing concentrations of DNA substrate for transformation experiments and found that both methods provided comparable results (Fig. 3). For both methods, transformation efficiencies were around 10^-6^ using 0.25 ng of *hu-sfgfp_aac* gDNA, then transformation frequencies increased by 10-fold with DNA concentration up to a transformation frequency around 10^-5^ for 2.5 ng of gDNA. However, for higher gDNA amounts (25 ng and 250 ng), the transformation frequencies seem to reach saturation and remained around 10^-4^. All together, these results demonstrate that detection of HU-sfGFP marker by flow cytometry constitutes a method as quantitative and specific as the classical antibiotic selection method to follow natural transformation in *A. baumannii.* The apramycin resistance cassette of the HU-sfGFP chromosomal marker was required here to benchmark the fluorescence-based method against the antibiotic selection method. However, our initial objective was to obtain a transformation marker that does not confer additional antibiotic resistance. We consequently engineered a HU-sfGFP chromosomal marker without an apramycin resistance cassette (Fig. S3A). We ensured that the new genetic construct was still compatible with bacterial growth (Fig. S3B) and that the fluorescence emitted was comparable to the *hu-sfgfp_aac* strain (Fig. S3C). Therefore, all following experiments were performed using the HU-sfGFP marker without any antibiotic resistance gene.

**Figure 3.**
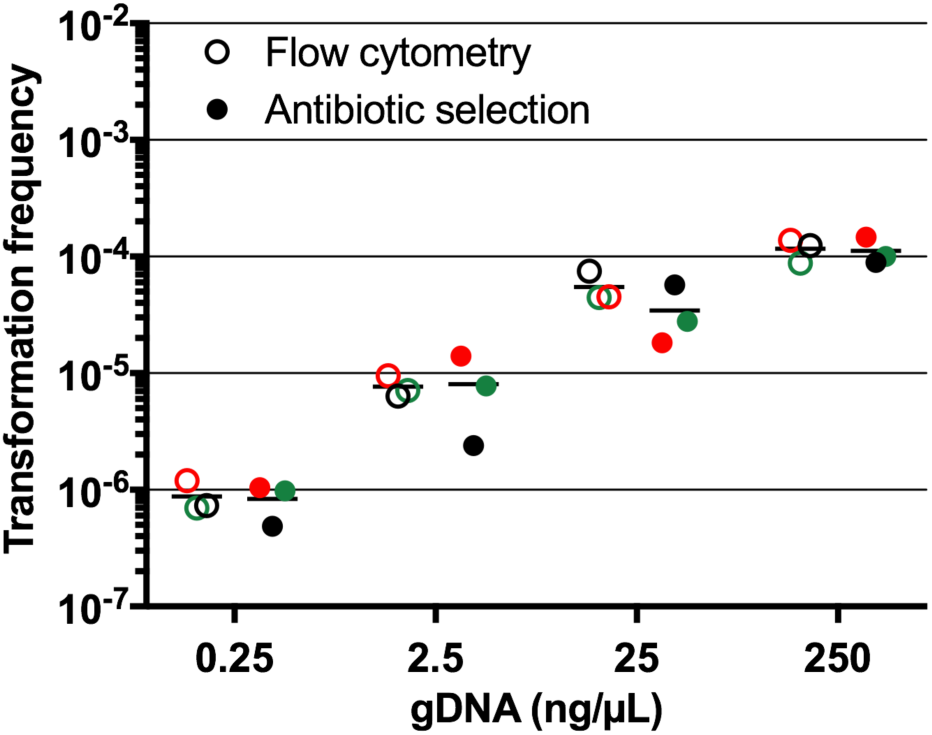
Comparison of flow cytometry sensitivity to antibiotic selection in detecting natural transformation with the *hu-sfgfp_aac marker*. Wild type *A. baumannii* AB5075 cells were transformed using increasing amounts of *hu-sfgfp_aac* genomic DNA (gDNA). DNA concentrations are expressed in nanogram of DNA per pL in the bacterial suspension used in transformation assay (abscissa). Transformation frequencies were determined both by flow cytometry (empty dots) and antibiotic selection (filled dots) for each gDNA concentration. For each method and DNA concentration, the three dots represent technical replicates. For a given gDNA concentration, dots of the same color indicates a sample tested by both methods. The horizontal lines represent the mean of the three measures.

### Improving parameters of natural transformation of *A. baumannii*

With the goal to detect natural transformation in MDR clinical isolates, we first took advantage of the fluorescent-based method to optimize conditions of natural transformation in *A. baumannii*. Because less than a ng of gDNA already results in frequencies higher than 10^-6^, we set this value as detection threshold of transformation. We first investigated how the type of transforming DNA would influence the transformation frequencies. *A. baumannii* strain AB5075 was subjected to transformation using the same mass of DNA (50 ng) carrying the HU-sfGFP marker flanked by 2 kb-long regions homologous to the targeted chromosomal locus, either as a linear PCR product, as an insertion in non-replicative plasmid (pASG-5) or as integrated at its locus in gDNA (Fig. 4). Transformation using gDNA resulted in transformation efficiencies between 10^-5^ and 10^-4^, comparable to the foregoing result (Fig. 3, 25 ng). However, transformation using a PCR product appeared less efficient with at least 10-fold fewer transformants than with gDNA. On the other hand, transformation frequencies using pASG-5 were significantly higher than using gDNA (mean of 1.5×10^-4^). As previously proposed for *A. baylyi* (30), the difference in transformation efficiency between gDNA and plasmid DNA (pASG- 5) may be explained by the number of copies of markers, much greater when using plasmid DNA than in gDNA. Taken this result into consideration, the following experiments were performed using pASG-5 as substrate of transformation. We tested first the impact of the bacterial culture and storage conditions prior to transformation assay and found no significant difference in transformation efficiencies after that bacteria were either grown on solid or liquid media or even stored at -80°C in conservation medium (Fig. S4). The optimal pH for natural transformation has been reported to be above 6.5 for *A. baylyi* ADP1 and of 7.5 in *A. baumannii* A118 (30, 31). We therefore investigated the effect of pH on natural transformation in strain AB5075. To this end, transformation medium was buffered in the range of 5.3 to 8.3 using potassium phosphate buffering system as used in Palmen *et al* (30) (Fig. 5A). We observed that between pH 5.3 and pH 6.1 the transformation levels were around 10^-4^. For pH greater than 6.36, the transformation frequencies decrease by 10-fold to decrease steadily down to 100-fold for pH above 7.36 to reach the lower detection limit. Based on the study of Traglia *et al.* performed on the *A. baumannii* strain A118, we also investigated the role of bovine serum albumin (BSA) on transformation efficiency to find no statistical difference on transformation level when a range from 0.25% to 1% BSA were added to the transformation medium (Fig. S5A). We also investigated the role of divalent cations (Ca^2+^, Mg^2+^ and Mn^2+^) in transformation but none of the cation, at the tested concentrations (1 and 5 mM for Ca^2+^and Mg^2+^, 0.5 mM for Mn^2+^), improved significantly the transformation levels (Fig. S5B). Finally, we tested the effect of agarose concentration in the transformation medium to observe that ranging from 0.25 to 1% of agarose, bacteria were increasingly transformable by up to one order of magnitude (Fig. 5B). However, above a concentration of 1% agarose, transformation reached a plateau.

**Figure 4.**
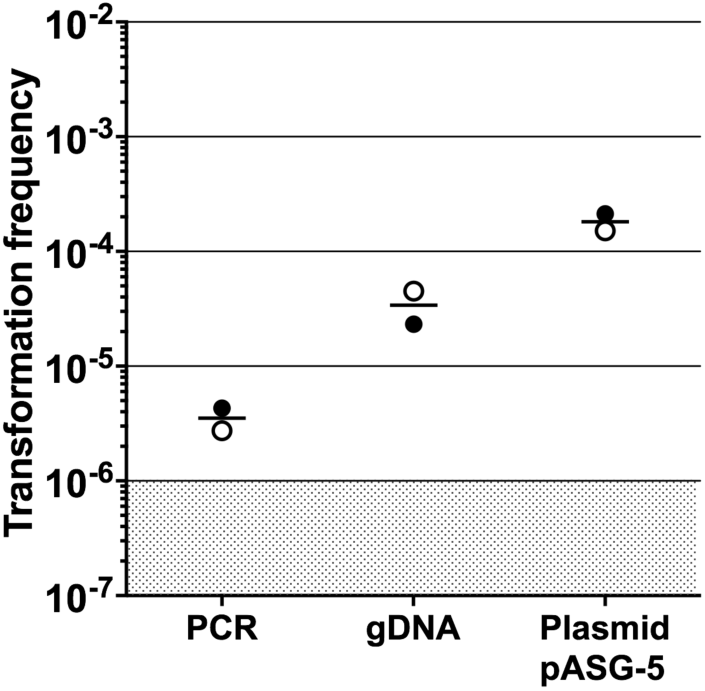
DNA forms and transformation efficiency. Comparison of transformation frequencies obtained with the *hu-sfgfp* marker on either a PCR fragment with 2 kb of flanking homology, genomic DNA or a non-replicative plasmid (pASG-5). The same mass of DNA (50 ng) was used for each DNA form. The experiment was performed twice in triplicates. Each dot of the same shape represents the mean of the triplicates obtained in the same independent experiment. The horizontal lines represent the mean of the two measurements. The limit of detection (10^-6^) is indicated by a shaded grey area.

**Figure 5.**
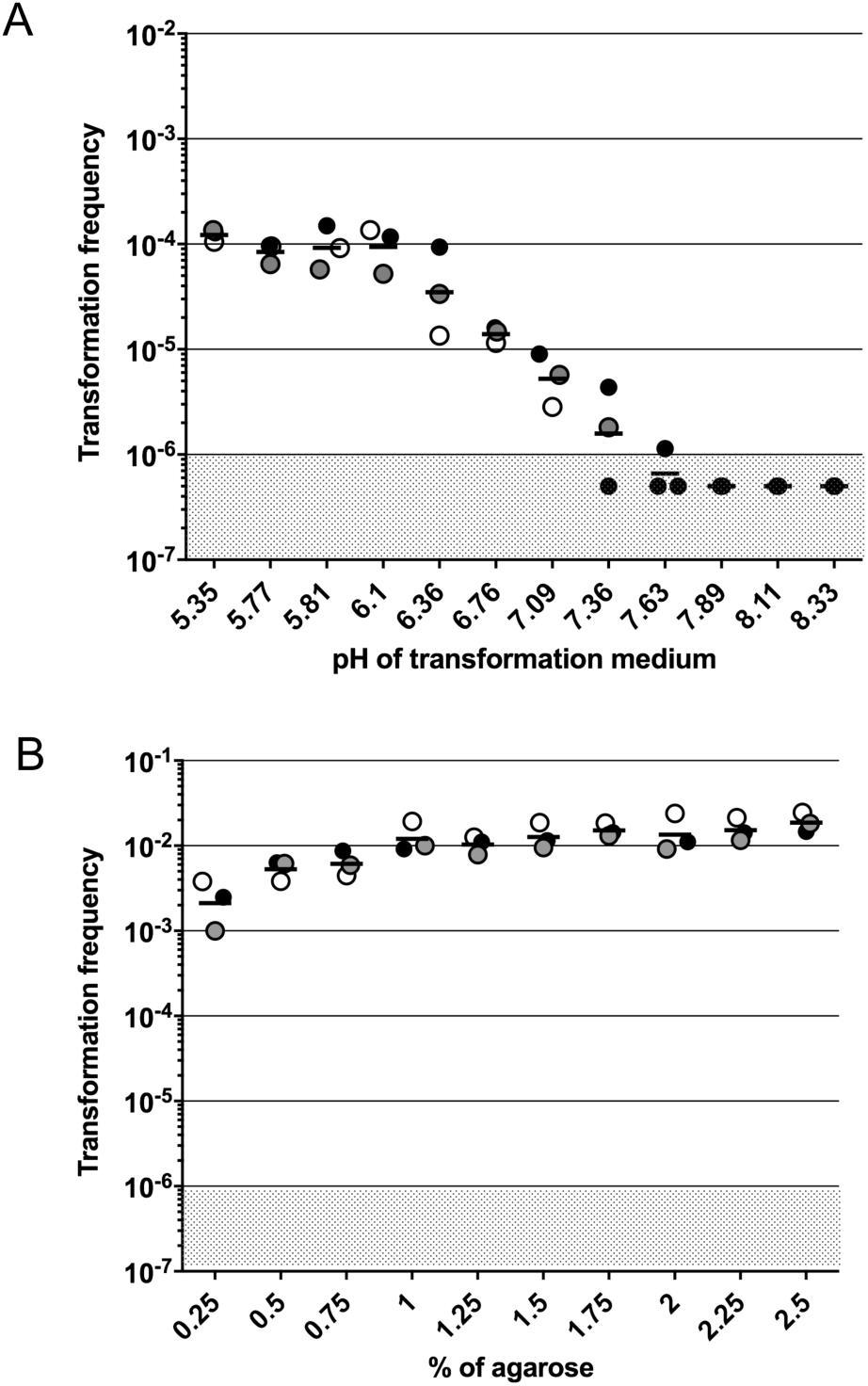
Parameters affecting transformability of *A. baumannii*. A. Effect of pH on transformation efficiencies. Bacteria were transformed using 0.5% agarose in transformation medium buffered using potassium phosphate with measured pH ranging from 5.35 to 8.33. B. Effect of increasing concentrations of agarose on transformation efficiencies (0.5% agarose was used in the foregoing experiments). Experiments were performed three times in triplicates. Each dot of the same shape represents the mean of the three replicates. The horizontal lines represent the mean of the three experiments. The limit of detection (10^-6^) is indicated by a shaded grey area.

Altogether, using flow cytometry, we refined the optimal conditions for transformation of *A. Baumannii* strain AB5075 (plasmid DNA, 2% agarose) and improved by 100-fold the transformation frequencies of this strain (comparison of Fig. 3 to Fig. 5B).

### Probing natural transformation in non-clinical and MDR clinical animal isolates

To further validate the cytometry-based method and to gain insight into the transformation ability of non-clinical strains isolated from wild animals, we sought to estimate the transformation ability of nine fully sequenced strains isolated from wild storks, some of them related to human clinical isolates (32). Taking advantage of avian strain genome availability (32), we confirmed that the HU gene and the flanking regions are highly conserved with at least 99,1% identity between the avian strains’ sequences and AB5075 strain sequence. Previously, five of these nine strains were found transformable in a qualitative transformation assay using the antibiotic-selection based method (32). All of the five strains were also found transformable using the fluorescence-based method but their ability to undergo transformation spreads over a range of three orders of magnitude (Fig. 6A). Remarkably, the fluorescence based-method allowed us to detect transformation in one isolate that was not identified as transformable using the antibiotic-selection method (32) (Fig. 6A, strain 280/1C). There is no clear correlation between transformation level and sequence identity of the transforming sequence given that strains that present the same sequence identity (99,4%, Fig. 6A) are either highly transformable (strains 29D2 and 86II/2C) or not transformable (strains 29R1 and 151/1C). Remarkably, strains presenting levels of sequence identity lower than the strain AYE (strains 29D2, 192/2C and 86II/2C) still presented comparable levels of transformation (Fig. 6A). Therefore, the transformation levels may reflect here the intrinsic ability of each strain to trigger competence and achieve transformation under these experimental conditions.

**Figure 6.**
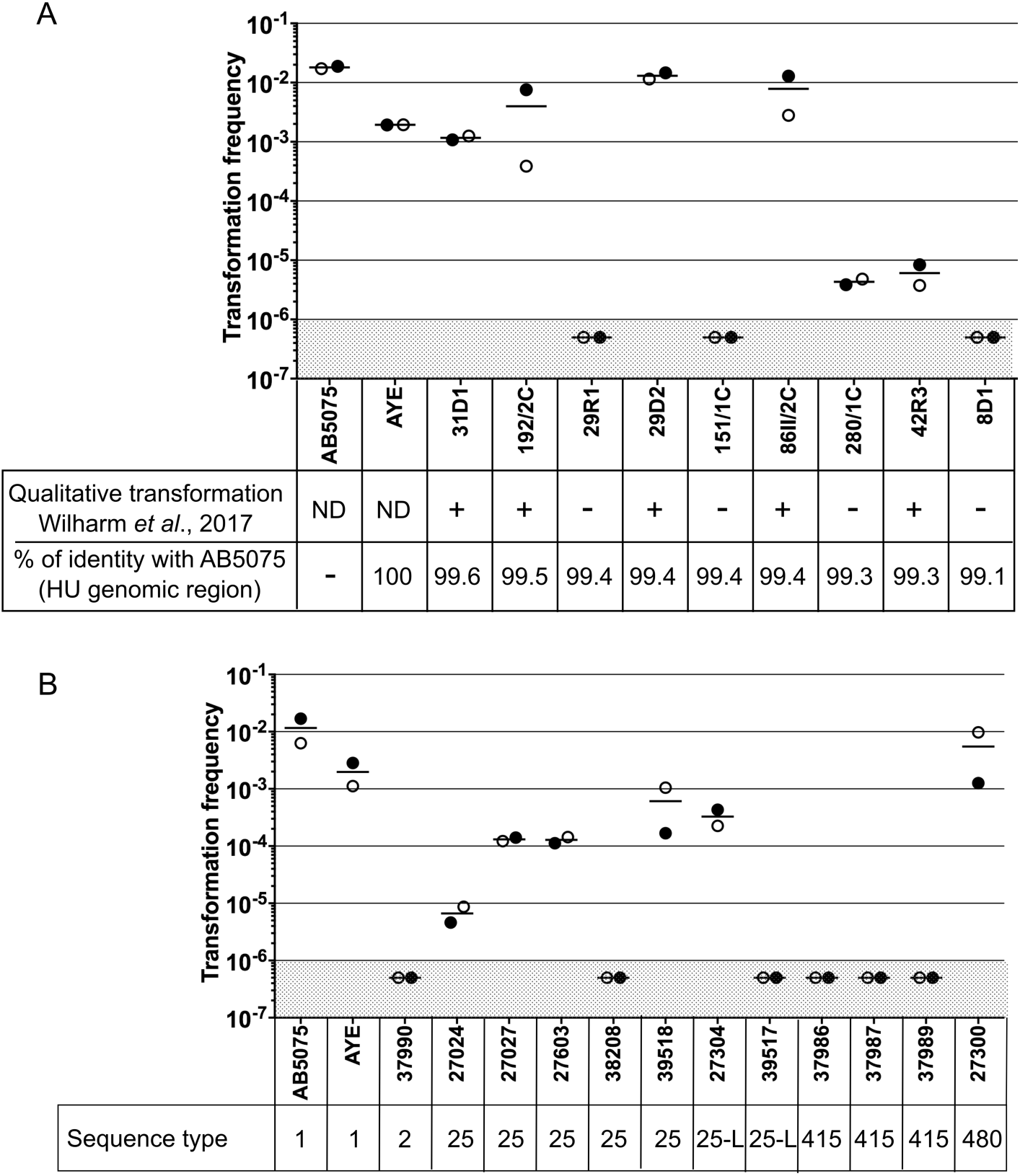
Determination of natural transformation frequencies of non-clinical and clinical *A. baumannii* isolates using flow cytometry. Wild animal (A) clinical animal isolates of *A. baumannii* (B) were subjected to transformation using pASG-5 and the optimized conditions developed for the strain. (A) Results of transformation ability obtained in a qualitative manner are reproduced from Wilharm et at., 2017 (ND: not done, “+” transformable, not transformable) (32). Percentage of sequence identity between HU region of strain AB5075 (4220 nucleotides long) with the recipient strain is indicated. (B) Sequence type (Pasteur scheme) of clinical isolates are mentionned (25-L standing for ST25-like). Transformability was determined twice independently, each dot represents the mean of three technical replicates. The horizontal lines represent the mean of the two experiments. The limit of detection (10^-6^) is indicated by a shaded grey area.

Taking advantage of the flow cytometry-based method and improved conditions for testing transformation, we also investigated the transformation profile of clinical MDR *A. baumannii* isolates of animal origin. We estimated the transformation frequencies of twelve *A. baumannii* isolates from diseased animals obtained through the French surveillance network for antimicrobial resistance in animal pathogens (Resapath). These isolates presented multiple antibiotic resistance including to carbapenems (33). For 66% of the isolates (8 out of 12) transformation could be detected, with transformation frequencies ranging from 10^-6^, the limit of detection, up to 10^-2^ (Fig. 6B). Interestingly, most isolates from the sequence type ST25 were found transformable contrarily to all the ST415 isolates. Altogether, these results on animal clinical isolates and non-clinical isolates demonstrate that the fluorescence-based method allows the detection of transformation events in *A. baumannii* regardless of the origin of the strain.

## DISCUSSION

*Acinetobacter baumannii* is now an established member of the growing list of bacteria capable of undergoing natural genetic transformation (13, 32). Together with genomic evidence of high recombination rates and multiple horizontal gene transfer events, natural transformation should be considered as a major contributor of antibiotic resistance genes acquisition in *A. baumannii*. To investigate natural transformation in clinical MDR isolates we developed here a fluorescence-based assay to detect and quantify transformation. To select an optimal marker of transformation, we engineered seven chromosomal markers consisting in C-terminal translational fusion of sfGFP with abundant proteins expressed from their original genetic environment. Interestingly, we obtained a Fis-sfGFP fusion protein, although a likely essential protein in AB5075 (34), but expression of this fusion did not confer the cells a fluorescence level distinguishable from autofluorescence (Fig. 1B, C and S1). All other translational fusions were highly expressed and emitted a fluorescence signal that could distinguish the corresponding strains from the wild-type strain (Fig. 1 and S1). Expectedly, the fluorescence intensity of the cells correlates with abundance of the sfGFP fusion protein (Fig. S1). Yet, protein folding or its location in the bacterial cell also influences the fluorescence intensity of a protein fusion. For instance, HU-sfGFP and HNS-sfGFP fusions are both predicted to be associated with the nucleoid and seemed to be expressed at similar level (Fig. S1). However, the HU-sfGFP fusion was brighter than the HNS one regardless of the growth phase (Fig. 1 and S2). While the HU-sfGFP signal appears distributed evenly in the nucleoid (Fig. 1B), the HNS-sfGFP signal was restricted to one to two discrete foci per cell. Both subcellular localizations observed in *A. baumannii* were reminiscent of localization patterns observed for HU and H-NS in *Escherichia coli* (35). Besides their use as fluorescent markers of natural transformation, the fusion proteins could be useful to further investigate the function of nucleoid-associated proteins in *A. baumannii*.

The HU-sfGFP marker combined with flow cytometry performed at least as well as the classical antibiotic selection to determine transformation frequencies (Fig. 2 and 3). We thus investigated the chemical and physical parameters susceptible to alter transformation efficiencies in *A. baumannii*. We tested the influence of divalent cations Mg^2+^, Ca^2+^ and Mn^2+^ on transformation efficiency to find only a mild effect of Ca^2+^ and Mn^2+^, as in *A. baylyi*, but not Mg^2+^. BSA was found to increase transformation by 2-fold in A118, yet the addition of this blood protein to the transformation medium did not affect transformation efficiency in AB5075 (Fig. S5A). With difference greater than two orders of magnitude, the parameter influencing transformation the most is pH (Fig. 5B). We found that transformation is more efficient at slightly acidic pH (5-6), which is opposite to results obtained in *A. baylyi* ADP1 for which transformation drops by 10 to 100-fold at pH below neutrality (30). This is a clear indication that even closely related species can undergo natural transformation under specific conditions. Transformation occurring at mildly acidic pH in AB5075 led us to hypothesize that competence may be triggered in several specific sites within a host. For instance, acidic pH are reminiscent of cutaneous pH, (slightly lower than 5 (36)), or parts of the gastro-intestinal tract, with pH of 5 to 7 in the colon (37). Noteworthy, these body sites contain large bacterial communities, including *A. baumannii* or other *Acinetobacter* species, that could offer substrates for natural transformation and foster acquisition of antibiotic resistance genes (4, 5).

We also found that increasing agarose concentration improved transformation (Fig. 5A). A simple explanation for the apparent agarose concentration-dependent increase of natural transformation could be that at high concentrations, agarose physically impairs bacterial movement and somehow increases the local DNA/bacterial cell ratio and therefore transformation efficiency. Another hypothesis could be that mechanochemical perception of type IV pilus retraction on agarose stimulates competence. Indeed, sensing pili retraction was involved in induction of virulence genes expression and holdfast synthesis respectively in *Pseudomonas aeruginosa* and *Vibrio cholera* (38, 39). Also, and intriguingly, agarose is a necessary component of semi-solid media required to natural transform *A. baumannii* or closely-related species (13, 32, 40). One hypothesis could be that the competent state for natural transformation is induced by the presence of agarose itself. Indeed, agarose, a polymer produced by seaweed, is made of repeating units of disaccharide (D-galactose and 3,6- anhydro-L-galactopyranose). Some soil or marine isolates of *Acinetobacter* sp. produce an agarase able to degrade agarose (41, 42). Similarly to degradation products of chitin that induce competence in *Vibrio cholera* (43), the degradation products of agarose could represent the trigger of competence in *A. baumannii*. The regulatory cascade that triggers competence in *A. baumannii* is currently unknown and its identification may provide clues about the inducing cues. Regardless of the inducing mechanism, the improved assay results in the highest transformation frequencies ever reported for *A. baumannii* which can be as high as 1% for strain AB5075. The combined improved assay and method of detection made possible the investigation of natural transformability on a panel of non-clinical strains and clinical isolates. We found that 66% of *A. baumannii* clinical isolates isolated from pets (8 out of 12) and non-clinical strains isolated from wild birds (6 out of 9) were transformable. Among transformable strains or isolates, transformation levels vary from 10^-2^ down to the detection limit (10^-6^). The variability of transformation efficiencies among *A. baumannii* isolates is reminiscent of transformation levels observed in *A. baumannii* human clinical isolates (13) but also among isolates of other transformable species such as *S. pneumoniae* or *Haemophilus influenzae* (44, 45). We ruled out the hypothesis that HU genomic region divergence prevents transformation.

However, although HU gene and protein sequences are highly conserved, the expression level of the HU gene in the recipient strain varies with some strains expressing less the HU gene in the condition tested resulting in cells presenting lower fluorescent signal (Fig. S6). Therefore, although the strain may be transformable, it is possible that transformants may escape detection. As discussed beforehand, another explanation for the undetectable transformability may be related to strain-specific difference in the conditions required for competence development. In this regard, the fact that most of the ST25 strains presented high level of transformation indicates that competence development is favored by the experimental conditions set for ST1 strains (AB5075 and AYE). These conditions may not be optimal for some strains and we cannot rule out that strains that appear non-transformable in our assay would someday reveal transformable under slightly different conditions. Indeed, a major limitation of our assay lies in its relative low sensitivity. Using our current equipment and settings, our assay does not allow us to conclude about the transformability of strains that would transform at frequencies below 10^-6^. Yet, the use of HU-sfGFP and flow cytometry offers, for the time, the possibility to phenotypically detect transformation events in clinical isolates of *A. baumannii* for which transformability could not have been otherwise investigated. Little is known about the conditions that foster horizontal gene transfer by transformation in *A. baumannii*. Our easy and rapid method now allows the study of the environmental conditions that may influence on rates of horizontal gene transfer, and this, using undomesticated strains. For instance, the method could be used to quantify horizontal gene transfer by natural transformation resulting from intra- or interspecific predatory behaviors (46, 47). The new HU-sfGFP would also constitute an excellent tool for the live-cell imaging of transformation in this species. It also opens investigations on horizontal gene transfer in infection models, experimentally testing the possibility that conditions specific to body sites, exposure to antibiotics or antiseptics may stimulate HGT.

### Materials and methods

#### Bacterial strains, typing and growth conditions

The bacterial strains or strains used in this study are listed in Table S1. Genotyping of the isolates was based on sequence type determination according to the multi-locus sequence typing proposed by (48). Unless specified, *Acinetobacter baumannii* isolates and strains were grown in lysogeny broth (LB) (Lennox). All experiments were performed at 37°C.

#### Construction of bacterial strains and plasmids

All the oligonucleotides used in this study for genetic modification are listed in supplementary Table S2. Plasmid pASG-1 was constructed by cloning chimeric PCR product from assembly of PCR1 (primers asg-2 and asg-4 on pKD-sfGFP obtained from Erwan Gueguen (Université de Lyon)) and PCR2 (primers Apr_Fw3 and asg-3 on plasmid carrying an apramycin resistance cassette) into pX5, a pMMB207 derivative to place the *sfgfp* gene under control of an artificial strong constitutive promoter. The plasmid pASG-5 was constructed by cloning a PCR performed on genomic DNA extracted from AB5075 *hu-sfGFP* strain using primers mlo-32 and mlo-35 into pJET1.2 following manufacturer instructions (CloneJET PCR cloning Kit, Thermofisher scientific). Plasmid pMHL-2 is a pGEM-t easy derivative in which an Apramycin resistance cassette and *sacB* counter-selection cassette have been cloned. All plasmid sequences are available upon request.

Gene disruptions were performed using overlap extension PCR to synthesize large chimeric DNA fragment carrying the selection/detection marker flanked by 2 kb fragments that are homologous to the insertion site. The oligonucleotides used for strain construction are listed in Table S2. Briefly, the sequence of the GFP superfolder (sfGFP) and the apramycin resistance cassette (*aac(3)IV*) were amplified from plasmid pASG-1 using primers mlo-28 and mlo-29. The DNA fragments allowing homologous recombination (2 kb upstream and downstream of the targeted locus) were obtained by PCR on genomic DNA of strain AB5075. In a second step, the fragments previously obtained were assembled by PCR. The PCRs were performed with a high-fidelity DNA polymerase (PrimeStarMax, Takara). Subsequently each PCR fragments were purified after migration on agarose gel followed by extraction as recommended by the manufacturer (QIAquick gel extraction kit, Qiagene). For fusion between nucleoproteins and sfGFP, a linker sequence which encodes the RGSGGEAAAKAGTS sequence between sfGFP and nucleoprotein has been added so as not to disturb the functioning of the proteins (25). All chimeric PCR products were independently introduced into the AB5075 wild type strain using natural transformation as described in the dedicated section.

To obtain a chromosomal marker without antibiotic cassette (*hu-sfgfp* strain), we first inserted a *sacB-aac* cassette (amplified from pMHL-2 plasmid) into the *hu* gene using overlap extension PCR. The resulting strain was subsequently transformed with a chimeric PCR product carrying a *hu-sfgfp* fusion without antibiotic determinant and counter-selected recombinants on minimal medium (M63) with 10% sucrose.

#### Fluorescence microscopy

Five hundred μl of culture were incubated 30 minutes at room temperature with formaldehyde (final concentration 3.7%), DAPI (final conc. 3 μM) and FM4-64 (final conc. 10 µg/mL). Three μl of the preparation were spotted on a poly-L-lysine coated coverslip. After adhesion of the bacterial cells, the coverslips were washed twice with Phosphate Buffer Saline solution (PBS). The specimens were then observed under a fluorescence microscope by an immersion objective for a final magnification of 1000X (EVOS FL, LifeTechnologies).

#### Flow cytometry

Bacterial suspensions were fixed with formaldehyde (final conc. 3.7%) and membrane stained with FM4-64 (final conc. 10 µg/mL) during 30 minutes at room temperature then washed twice with PBS and further diluted in PBS to obtain about 10^6^ cells/mL. Attune Acoustic Focusing Cytometer (Life Technologies) was used for all flow cytometry acquisitions. Samples were run at a collection rate of 25 μL/min, and fluorescence emission was detected using a 530/30 bandpass filter for GFP fluorescence and 640 longpass filter for FM4-64 fluorescence. To avoid aggregates a maximum concentration of 10^6^/mL FM4-64–positive particles were analyzed. Flow cytometry data were analyzed using the Attune software. Green fluorescence was analyzed on a minimum of 20 millions of FM4-64–positive particles.

#### Transformation assay

The following method was adapted from reference (13). Overnight cultures at 37°C in LB liquid medium are diluted in Phosphate Buffer Saline (PBS) to obtain 10^7^ CFU/mL. Then 10 µL of bacterial suspension is mixed with 10 µL of substrate DNA (at 100 ng/µL for genomic DNA) and 2.5 µL of this bacterial/DNA suspension is then stabbed into 1mL of freshly prepared motility medium (5 g/L Agarose, 5 g/L Tryptone and 2.5 g/L NaCl) in 2 mL Eppendorf tubes. After 20 h at 37°C, 200 µL of Phosphate Buffer Saline (PBS) is added to the tube and bacteria are resuspended by vigorous vortexing. The transformants are then selected by plating on selective agar media (apramycin 30 µg/µL) or their fluorescence measured by flow cytometry. Determination of transformation frequencies is then obtained by calculating the ratio of the number of transformant CFUs (apramycin resistant or fluorescent) on the total number of CFU (antibiotic sensitive or non-fluorescent).

All the transformation assays were performed twice or three times in triplicates. To ensure transparency and as proposed (49) the means of the replicates are shown as a single independent data point and all the independent data points are plotted.

#### Funding Information

This work was supported by the LABEX ECOFECT (ANR-11-LABX-0048) of Université de Lyon, within the program “Investissements d’Avenir” (ANR-11-IDEX-0007) operated by the French National Research Agency (ANR). ASG and MHL were also supported by *Programme Jeune Chercheur* from VetAgro Sup. AL and MH were supported by the French Agency for Food, Environmental and Occupational Health & Safety (ANSES). G.W. received funding from the Deutsche Forschungsgemeinschaft (DFG) within FOR 2251 (WI 3272/3-1).

## References

1. Dijkshoorn L, Nemec A, Seifert H. 2007. An increasing threat in hospitals: multidrug-resistant Acinetobacter baumannii. Nat Rev Microbiol 5:939–951.

2. Zordan S, Prenger-Berninghoff E, Weiss R, van der Reijden T, van den Broek P, Baljer G, Dijkshoorn L. 2011. Multidrug-resistant Acinetobacter baumannii in veterinary clinics, Germany. Emerg Infect Dis 17:1751–1754.

3. Eveillard M, Kempf M, Belmonte O, Pailhoriès H, Joly-Guillou M-L. 2013. Reservoirs of Acinetobacter baumannii outside the hospital and potential involvement in emerging human community-acquired infections. Int J Infect Dis IJID Off Publ Int Soc Infect Dis 17:e802–5.

4. Zeana C, Larson E, Sahni J, Bayuga SJ, Wu F, Della-Latta P. 2003. The Epidemiology of Multidrug-Resistant <span class=“italic”>Acinetobacter Baumannii</span> Does the Community Represent a Reservoir? Infect Control Amp Hosp Epidemiol 24:275–279.

5. Dijkshoorn L, Aken E van, Shunburne L, Reijden TJK van der, Bernards AT, Nemec A, Towner KJ. 2005. Prevalence of Acinetobacter baumannii and other Acinetobacter spp. in faecal samples from non-hospitalised individuals. Clin Microbiol Infect 11:329–332.

6. Pailhoriès H, Kempf M, Belmonte O, Joly-Guillou M-L, Eveillard M. 2016. First case of OXA-24-producing Acinetobacter baumannii in cattle from Reunion Island, France. Int J Antimicrob Agents 48:763–764.

7. Hérivaux A, Pailhoriès H, Quinqueneau C, Lemarié C, Joly-Guillou M-L, Ruvoen N, Eveillard M, Kempf M. First report of carbapenemase-producing Acinetobacter baumannii carriage in pets from the community in France. Int J Antimicrob Agents.

8. European Centre for Disease Prevent (ECDC)ion and Control. 2017. Antimicrobial resistance surveillance in Europe 2016. Annual Report of the European Antimicrobial Resistance Surveillance Network (EARS-Net). ECDC, Stockholm.

9. World Health Organization. 2017. Global antimicrobial resistance surveillance system (GLASS) report: early implementation 2016-2017. Geneva.

10. Fournier P-E, Vallenet D, Barbe V, Audic S, Ogata H, Poirel L, Richet H, Robert C, Mangenot S, Abergel C, Nordmann P, Weissenbach J, Raoult D, Claverie J-M. 2006. Comparative genomics of multidrug resistance in Acinetobacter baumannii. PLoS Genet 2:e7. 1.

11. Wright MS, Haft DH, Harkins DM, Perez F, Hujer KM, Bajaksouzian S, Benard MF, Jacobs MR, Bonomo RA, Adams MD. 2014. New insights into dissemination and variation of the health care-associated pathogen Acinetobacter baumannii from genomic analysis. mBio 5:e00963–13.

12. Snitkin ES, Zelazny AM, Montero CI, Stock F, Mijares L, NISC Comparative Sequence Program, Murray PR, Segre JA. 2011. Genome-wide recombination drives diversification of epidemic strains of Acinetobacter baumannii. Proc Natl Acad Sci 108:13758–13763.

13. Wilharm G, Piesker J, Laue M, Skiebe E. 2013. DNA uptake by the nosocomial pathogen Acinetobacter baumannii occurs during movement along wet surfaces. J Bacteriol 195:4146–4153.

14. Touchon M, Cury J, Yoon E-J, Krizova L, Cerqueira GC, Murphy C, Feldgarden M, Wortman J, Clermont D, Lambert T, Grillot-Courvalin C, Nemec A, Courvalin P, Rocha EPC. 2014. The genomic diversification of the whole Acinetobacter genus: origins, mechanisms, and consequences. Genome Biol Evol 6:2866–2882.

15. Harding CM, Tracy EN, Carruthers MD, Rather PN, Actis LA, Munson RS. 2013. Acinetobacter baumannii strain M2 produces type IV pili which play a role in natural transformation and twitching motility but not surface-associated motility. mBio 4:e00360–13.

16. Bonnin RA, Poirel L, Nordmann P. 2012. AbaR-type transposon structures in Acinetobacter baumannii. J Antimicrob Chemother 67:234–236.

17. Krizova L, Dijkshoorn L, Nemec A. 2011. Diversity and evolution of AbaR genomic resistance islands in Acinetobacter baumannii strains of European clone I. Antimicrob Agents Chemother 55:3201–3206.

18. Kim DH, Ko KS. 2015. AbaR-type genomic islands in non-baumannii Acinetobacter species isolates from South Korea. Antimicrob Agents Chemother 59:5824–5826.

19. Johnston C, Martin B, Fichant G, Polard P, Claverys J-P. 2014. Bacterial transformation: distribution, shared mechanisms and divergent control. Nat Rev Microbiol 12:181–196.

20. Domingues S, Harms K, Fricke WF, Johnsen P al J, da Silva GJ, Nielsen KM. 2012. Natural transformation facilitates transfer of transposons, integrons and gene cassettes between bacterial species. PLoS Pathog 8:e1002837.

21. Seitz P, Blokesch M. 2013. Cues and regulatory pathways involved in natural competence and transformation in pathogenic and environmental Gram-negative bacteria. FEMS Microbiol Rev 37:336–363.

22. Dorer MS, Fero J, Salama NR. 2010. DNA Damage Triggers Genetic Exchange in Helicobacter pylori. PLOS Pathog 6:e1001026.

23. Charpentier X, Kay E, Schneider D, Shuman HA. 2011. Antibiotics and UV radiation induce competence for natural transformation in Legionella pneumophila. J Bacteriol 193:1114–1121.

24. Prudhomme M, Attaiech L, Sanchez G, Martin B, Claverys J-P. 2006. Antibiotic stress induces genetic transformability in the human pathogen Streptococcus pneumoniae. Science 313:89–92.

25. Kjos M, Aprianto R, Fernandes VE, Andrew PW, van Strijp JAG, Nijland R, Veening J-W. 2015. Bright fluorescent Streptococcus pneumoniae for live-cell imaging of host-pathogen interactions. J Bacteriol 197:807–818.

26. Dillon SC, Dorman CJ. 2010. Bacterial nucleoid-associated proteins, nucleoid structure and gene expression. Nat Rev Microbiol 8:185–195.

27. Soares NC, Cabral MP, Parreira JR, Gayoso C, Barba MJ, Bou G. 2009. 2-DE analysis indicates that Acinetobacter baumannii displays a robust and versatile metabolism. Proteome Sci 7:37.

28. Fay A, Glickman MS. 2014. An essential nonredundant role for mycobacterial DnaK in native protein folding. PLoS Genet 10:e1004516.

29. Lewis PJ, Thaker SD, Errington J. 2000. Compartmentalization of transcription and translation in Bacillus subtilis. EMBO J 19:710–718.

30. Palmen R, Vosman B, Buijsman P, Breek CK, Hellingwerf KJ. 1993. Physiological characterization of natural transformation in Acinetobacter calcoaceticus. J Gen Microbiol 139:295–305.

31. Rochelle PA, Day MJ, Fry JC. 1988. Occurrence, Transfer and Mobilization in Epilithic Strains of Acinetobacter of Mercury-resistance Plasmids Capable of Transformation. Microbiology 134:2933–2941.

32. Wilharm G, Skiebe E, Higgins PG, Poppel MT, Blaschke U, Leser S, Heider C, Heindorf M, Brauner P, Jäckel U, Böhland K, Cuny C, Łopińska A, Kaminski P, Kasprzak M, Bochenski M, Ciebiera O, Tobółka M, Żołnierowicz KM, Siekiera J, Seifert H, Gagné S, Salcedo SP, Kaatz M, Layer F, Bender JK, Fuchs S, Semmler T, Pfeifer Y, Jerzak L. Relatedness of wildlife and livestock avian isolates of the nosocomial pathogen Acinetobacter baumannii to lineages spread in hospitals worldwide. Environ Microbiol 19:4349–4364.

33. Lupo A, Châtre P, Ponsin C, Saras E, Jean-Boulouis H, Keck N, Haenni M, Madec J-Y. 2016. Clonal Spread of bla _OXA-23_ ST25 Acinetobacter baumannii in Companion Animals in France. Antimicrob Agents Chemother AAC.01881-16.

34. Gallagher LA, Ramage E, Weiss EJ, Radey M, Hayden HS, Held KG, Huse HK, Zurawski DV, Brittnacher MJ, Manoil C. 2015. Resources for Genetic and Genomic Analysis of Emerging Pathogen Acinetobacter baumannii. J Bacteriol 197:2027–2035.

35. Wang W, Li G-W, Chen C, Xie XS, Zhuang X. 2011. Chromosome organization by a nucleoid-associated protein in live bacteria. Science 333:1445–1449.

36. Matousek JL, Campbell KL. 2002. A comparative review of cutaneous pH. Vet Dermatol 13:293–300. 1.

37. Cook MT, Tzortzis G, Charalampopoulos D, Khutoryanskiy VV. 2012. Microencapsulation of probiotics for gastrointestinal delivery. J Controlled Release 162:56–67.

38. Persat A, Inclan YF, Engel JN, Stone HA, Gitai Z. 2015. Type IV pili mechanochemically regulate virulence factors in Pseudomonas aeruginosa. Proc Natl Acad Sci 112:7563–7568.

39. Ellison CK, Kan J, Dillard RS, Kysela DT, Ducret A, Berne C, Hampton CM, Ke Z, Wright ER, Biais N, Dalia AB, Brun YV. 2017. Obstruction of pilus retraction stimulates bacterial surface sensing. Science 358:535–538.

40. Harding CM, Tracy EN, Carruthers MD, Rather PN, Actis LA, Munson RS. 2013. Acinetobacter baumannii Strain M2 Produces Type IV Pili Which Play a Role in Natural Transformation and Twitching Motility but Not Surface-Associated Motility. mBio 4.

41. Lakshmikanth M, Manohar S, Lalitha J. 2009. Purification and characterization of β-agarase from agar-liquefying soil bacterium, Acinetobacter sp., AG LSL-1. Process Biochem 44:999–1003.

42. Leema Roseline T, Sachindra NM. 2016. Characterization of extracellular agarase production by Acinetobacter junii PS12B, isolated from marine sediments. Biocatal Agric Biotechnol 6:219–226.

43. Meibom KL, Blokesch M, Dolganov NA, Wu C-Y, Schoolnik GK. 2005. Chitin Induces Natural Competence in Vibrio cholerae. Science 310:1824–1827.

44. Evans BA, Rozen DE. 2013. Significant variation in transformation frequency in Streptococcus pneumoniae. ISME J 7:791–799.

45. Maughan H, Redfield RJ. 2009. Extensive variation in natural competence in Haemophilus influenzae. Evol Int J Org Evol 63:1852–1866.

46. Cooper RM, Tsimring L, Hasty J. 2017. Inter-species population dynamics enhance microbial horizontal gene transfer and spread of antibiotic resistance. eLife 6:e25950.

47. Borgeaud S, Metzger LC, Scrignari T, Blokesch M. 2015. The type VI secretion system of Vibrio cholerae fosters horizontal gene transfer. Science 347:63–67.

48. Diancourt L, Passet V, Nemec A, Dijkshoorn L, Brisse S. 2010. The Population Structure of Acinetobacter baumannii: Expanding Multiresistant Clones from an Ancestral Susceptible Genetic Pool. PLOS ONE 5:e10034.

49. Vaux DL. 2012. Research methods: Know when your numbers are significant. Nature 492:180–181.

